# Tissue-specific downregulation of *EDTP* suppresses polyglutamine protein aggregates and extends lifespan in *Drosophila*

**DOI:** 10.1101/130559

**Authors:** Chengfeng Xiao, Shuang Qiu, R Meldrum Robertson, Laurent Seroude

## Abstract

*Drosophila* egg-derived tyrosine phosphatase (EDTP, also called JUMPY) is a lipid phosphatase essential in oogenesis and muscle function in the adult stage. Although mammalian JUMPY negatively regulates autophagy, loss-of-JUMPY causes muscle dysfunction and is associated with a rare genetic disorder called centronuclear myopathy. Here we show that tissue-specific downregulation of EDTP in *Drosophila* non-muscle tissues, particularly glial cells, suppresses the expression of polyglutamine (polyQ) protein aggregates in the same cells and improves survival. Additionally, tissue-specific downregulation of EDTP in glial cells or motoneurons extends lifespan. We demonstrate an approach to fine-tune the expression of a disease-associated gene *EDTP* for the removal of polyQ protein aggregate and lifespan extension in *Drosophila.*

## Introduction

Egg-derived tyrosine phosphatase (EDTP, also called JUMPY) is a lipid phosphatase that removes 3-position phosphate at the inositol ring of phosphatidylinositol 3-phosphate (PtdIns3P) and phosphatidylinositol (3, 5)-bi-phosphate (PtdIns(3,5)P_2_)^1^. EDTP has an opposite function to Vps34, a sole class III phosphoinositide 3-kinase^2,3^, in the regulation of the PtdIns3P pool.

The most interesting characteristic of EDTP expression in *Drosophila* is that there are two peaks, one at oogenesis^4,5^, and the other at the adult stage^6^. The transcription of *mJUMPY,* a mouse homolog to *EDTP,* also shows a peak at day five of differentiation of C2C12 myoblasts, followed by a decline^1^. In addition, human JUMPY is detectable in all tested tissues at the ages of 19 - 69 years^1^. Another expression characteristic is that - human *JUMPY* is ubiquitously expressed and abundant in skeletal muscle^1^. Such an expression pattern is highly coincident between *JUMPY* and *Vps34^7^,* indicating a tight regulation of PtdIns3P levels. The decline of EDTP between early embryogenesis and the young adult stage is accompanied with *Drosophila* metamorphosis, a process requiring extensive autophagy and apoptosis for histolysis^8^. This observation suggests a role for EDTP in the regulation of autophagy. This is indeed supported by the findings that PtdIns3P stimulates autophagy in human HT-29 cells^9^, that mJUMPY negatively controls autophagosome formation and maturation in mammalian cells^10^, and that EDTP/JUMPY inhibitors, AUTEN-67 and AUTEN-99, activate autophagy in human cell lines and mouse tissues^11,12^.

Despite the negative regulation of autophagy, a *Drosophila* null *EDTP* mutant is lethal at embryogenesis or in the first instar, and germline clones with a null *EDTP* allele fail to produce mature oocytes^5^. Homozygous flies carrying a hypomorphic allele of *EDTP* are short-lived with impaired motor functions and reduced fecundity^13^. Additionally, muscles of the mJUMPY-deficiency mouse have decreased force production, prolonged relaxation and exacerbated fatigue^14^. Human JUMPY missense variant (R336Q) has a link to centronuclear myopathy, a rare genetic disorder with muscle weakness and wasting^1^. Therefore, the function of autophagy initiation is likely overwhelmed by the lethality or disease-causing effects of loss-of-JUMPY ubiquitously or in the muscles.

There are advantages of autophagy in degrading and recycling disrupted organelles, long-lived proteins and denatured protein aggregates^9,15^. A strategy to maximize the potentially beneficial effects of EDTP is to manipulate its expression in selected tissues while keeping EDTP unaffected in the muscles. This seems to be feasible by using the *Drosophila* Gal4/UAS expression system^16^. We therefore hypothesized that selective downregulation of EDTP in the central nervous system removes protein aggregates and extends lifespan in *Drosophila.*

In the current study, we performed several sets of experiments to examine the effects of selective downregulation of EDTP on the removal of protein aggregates and lifespan extension in *Drosophila*. We first demonstrate that heterozygous EDTP mutation improves survival after prolonged anoxia exposure, a stress which induces autophagy^17^. We next show that RNAi knockdown of EDTP in glial cells suppresses the expression of polyglutamine (polyQ) protein aggregates in the same cells. Finally, RNAi knockdown of EDTP in glial cells or motoneurons extends lifespan in *Drosophila.*

## Results

### Improved survival after prolonged anoxia in *EDTP* mutant

Heterozygous flies of DJ694, an *EDTP* mutant^6,13^, were exposed to a prolonged anoxia of six hours at the ages of 7-8 days. DJ694/+ males showed overall improved survival (median 43 days, n = 167) compared with their sibling controls w^1118^ flies (median 8 days, n = 163) (P = 0.0033, Mantel-Cox test) (Fig. 1a). Female DJ694/+ flies also showed better survival (median 70.5 days, n = 150) than controls (median 18 days, n = 148) (P = 0.0001, Mantel-Cox test) (Fig. 1b). Without anoxia exposure, survival was the same between DJ694/+ males (median 76 days, n = 102) and w^1118^ males (median 76 days, n = 96) (P = 0.9452, Mantel-Cox test) (Fig. 1c). Survival was also the same between DJ694/+ females (median 81 days, n = 107) and w^1118^ females (median 81 days, n = 97) (P = 0.8688, Mantel-Cox test) (Fig. 1d). Heterozygous *EDTP* mutants displayed improved survival to a 6-h anoxia.

**Fig. 1.**
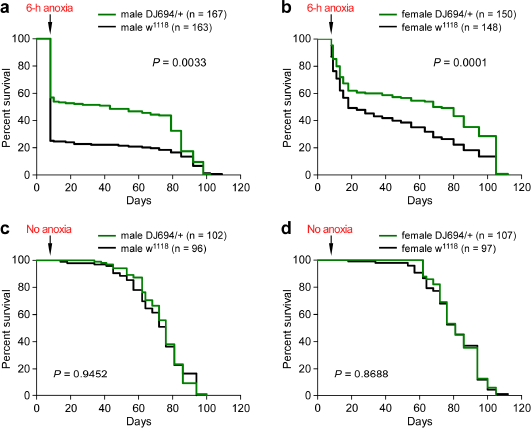
Improved survival after prolonged anoxia in *EDAP* mutant. (a) Survival of male flies after an exposure to 6-h anoxia. Anoxia was applied to flies at the ages of 7-8 days. DJ694/+ and w1118 flies were siblings prepared by two consecutive crosses. Numbers of flies (n) are indicated. DJ694 is an EDTP mutant line^6^. (b) Survival of female flies with an exposure to 6-h anoxia. Flies were treated the same way as males. (c) Survival of male flies without anoxia exposure. (d) Survival of female flies without anoxia exposure.

### Polyglutamine protein aggregates expressed in glial cells

Extreme hypoxia induces autophagy in mammalian and human cells^17^. *Drosophila* EDTP might have functions similar to its mammalian counterpart in the negative regulation of autophagy. To support this proposal, a *Drosophila* model with protein aggregates expressed in the central nervous system is desired. We directed the expression of a DNA construct, UAS-Httex1-Q72-eGFP^18^, in glial cells with repo-Gal4^19,20^. This UAS line has been successfully used in the expression of polyQ protein aggregates in the compound eyes^18^.

PolyQ protein aggregates were clearly observed in the brain of UAS-Httex1-Q72-eGFP/+;;repo-Gal4/+ fly (Fig. S1). No polyQ protein aggregate was observed in the brains of control flies (UAS-Httex1-Q72-eGFP/+ and repo-Gal4/+). Expression of heat shock protein 70^21^, a soluble molecular chaperone, did not form protein aggregates in glial cells in repo-Gal4/UAS-hsp70-myc fly. Also, expression of mCD8-GFP, a membrane-anchored protein, resulted in no protein aggregates in glial cells in repo-Gal4/20×UAS-IVS-mCD8::GFP fly. The polyQ protein aggregates expressed in glial cells were used as an indicator to examine the effects of *EDTP* downregulation.

### Suppression of polyQ protein aggregates by RNAi knockdown of ***EDTP*** in glial cells

PolyQ protein aggregates were observed in the brains of UAS-Httex1-Q72-eGFP/+;;repo-Gal4/+ flies. At day 1, 5, 10 and 19, the aggregates were seen widely spread in the brain. Large aggregates were observed in the surface region, median septum and connections between middle brain and optic lobes. Relatively small aggregates were diffused throughout the brain. Few flies survived more than 19 days (Fig. 2a).

**Figure 2.**
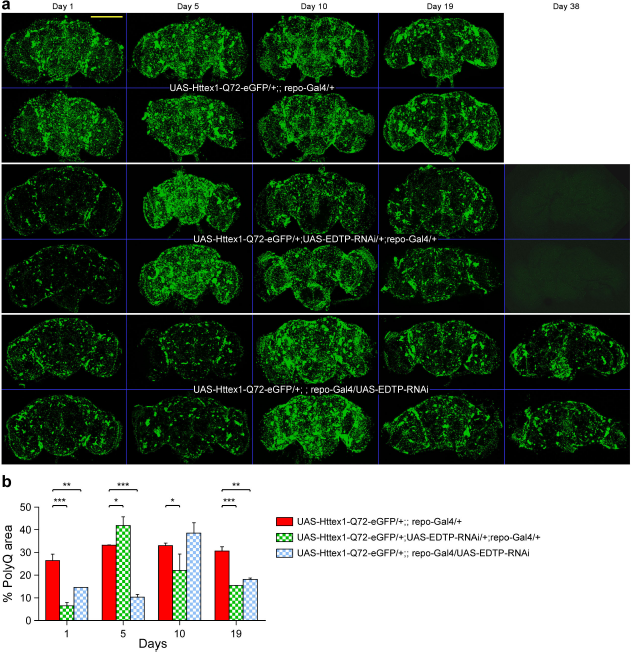
Suppression of polyQ protein aggregates by *EDTP* downregulation. (a) Expression of polyQ protein aggregates in glial cells in different flies. Shown are the brains of male flies (n = 2) at the ages of 1, 5, 10, 19 and 38 days. The genotype designations are embedded in the images. The images at day 38 for UAS-Httex1-Q72-eGFP/+; repo-Gal4/+ flies are unavailable, because few flies survive longer than 19 days. PolyQ protein aggregates can not be detected in the brains of UAS-Httex1-Q72-eGFP/+; UAS-EDTP-RNAi/+; repo-Gal4/+ flies. Bar 200 mm. (b) Analysis of the percent area for polyQ protein aggregates (% PolyQ area) between flies. Fly ages are indicated. * P < 0.05; ** P < 0.01; *** P < 0.001 by two-way ANOVA with Bonferroni post-tests.

We examined the effect of *EDTP* downregulation on the expression of polyQ protein aggregates. *EDTP* knockdown was performed by using an RNAi line carrying dsRNA to *EDTP* on the second chromosome (BDSC #41633). This RNAi line has been used to knockdown EDTP in fat bodies in larvae^11^. PolyQ protein aggregates were seen in the brains at day 1, 5, 10 and 19 of UAS-Httex1-Q72-eGFP/+; UAS-EDTP-RNAi/+; repo-Gal4/+ flies. These flies lived up to 38 days, which was a 2-fold increase in maximal survival compared with flies without *EDTP* downregulation. Strikingly, polyQ protein aggregates were invisible at day 38 in flies with *EDTP* knockdown. The disappearance of polyQ protein aggregates was observed at as early as day 30 (Fig. S2).

Using another RNAi line, which carried a dsRNA to *EDTP* on the third chromosome, we observed that polyQ protein aggregates were present in the brains at day 1, 5, 10, 19 and 38. Again, flies with *EDTP* downregulation had roughly a 2-fold increase in maximal survival relative to controls. However, polyQ protein aggregates were clearly seen in the brains at day 38. Thus, complete disappearance of polyQ protein aggregates was observed by using one RNAi but not another (Fig. 2a).

At day 1, the percent area for polyQ protein aggregates (*%* polyQ area) was 26.5 *%* in UAS-Httex1-Q72-eGFP/+;;repo-Gal4/+ flies. The levels (26.5 - 33.2 *%*) remained relatively stable throughout the tested ages (Fig. 2b). *%* polyQ area was significantly reduced at day 1, 10 and 19 but not day 5 in UAS-Httex1-Q72-eGFP/+; UAS-EDTP-RNAi/+; repo-Gal4/+ flies compared respectively with the same-aged controls (Two-way ANOVA with Bonferroni post-tests) (Fig. 2b). At day 5, however, *%* polyQ area in flies with *EDTP* knockdown was even higher than that in control (P < 0.05, Two-way ANOVA with Bonferroni post-tests). Thus, *EDTP* downregulation suppressed polyQ protein aggregates at most tested ages, and reshaped the expression dynamics of polyQ protein aggregates.

*%* polyQ area was reduced at day 1,5 and 19 but not day 10 in UAS-Httex1-Q72-eGFP/+;;repo-Gal4/UAS-EDTP-RNAi flies compared with respective controls (Two-way ANOVA with Bonferroni post-tests) (Fig. 2b). Similarly, *EDTP* downregulation by a different RNAi suppressed polyQ protein aggregates and reshaped their expression dynamics.

Flies with two RNAi (UAS-Httex1-Q72-eGFP/+;UAS-EDTP-RNAi/+;repo-Gal4/UAS-EDTP-RNAi) survived up to 38 days. PolyQ protein aggregates were observable at day 1, 10, 19 and 38 (Fig. S3). Thus, we did not observe better suppression of polyQ protein aggregates using two RNAi rather than one.

### RNAi knockdown of ***EDTP*** in glial cells improved survival with polyQ protein aggregates

We examined the survival of flies with polyQ protein aggregates expressed in glial cells. The survival of UAS-Httex1-Q72-eGFP/+;;repo-Gal4/+ flies (median 15 days, n = 44) was remarkably shorter than repo-Gal4/+ flies (median 76 days, n = 37) (P < 0.0001, Mantel-Cox test) or UAS-Httex1-Q72-eGFP/+ flies (median 84 days, n = 62) (P < 0.0001, Mantel-Cox test) (Fig. 3a). Most UAS-Httex1-Q72-eGFP/+;;repo-Gal4/+ flies died within 20 days. This was consistent with the observation that few flies with the ages greater than 19 days were available for imaging.

**Figure 3.**
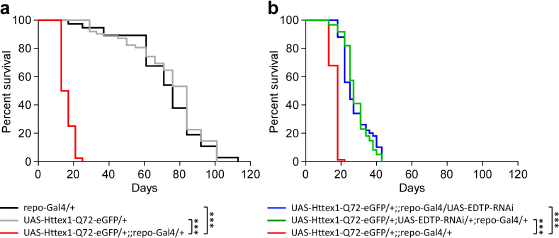
Improved survival with polyQ protein aggregates by *EDTP* downregulation. (a) Expression of
polyQ protein aggregates in glial cells shortened the lifespan. Survival of UAS-Httex1-Q72-eGFP/+;;repo-Gal4/+ flies (red) was markedly decreased compared with repo-Gal4/+ (black) or UAS-Httex1-Q72-eGFP/+ (grey). ***, P < 0.0001, Mantel-Cox test. Male flies were examined. (b) EDTP downregulation in glial cells improved survival with polyQ protein aggregates. Survival of UAS-Httex1-Q72-eGFP/+;; repo-Gal4/UAS-EDTP-RNAi flies (blue) or UAS-Httex1-Q72-eGFP/+; UAS-EDTP-RNAi/+; repo-Gal4/+ flies (green) was improved compared with UAS-Httex1-Q72-eGFP/+;; repo-Gal4/+ flies (red). Experiments with two independent RNAi lines showed similar results. ***, P < 0.0001, Mantel-Cox test.

The survivorship was examined in flies with the expression of polyQ protein aggregates and simultaneous RNAi knockdown of *EDTP* in glial cells. The survival of UAS-Httex1-Q72-eGFP/+; repo-Gal4/UAS-EDTP-RNAi (median 25 days, n = 50) was longer than that of UAS-Httex1-Q72-eGFP/+;repo-Gal4/+ flies (median 18 days, n = 84) (P < 0.0001, Mantel-Cox test). The survival of UAS-Httex1-Q72-eGFP/+;; UAS-EDTP-RNAi/+; repo-Gal4/+ (median 27 days, n = 61) was also longer than that of UAS-Httex1-Q72-eGFP/+;repo-Gal4/+ flies (P < 0.0001, Mantel-Cox test) (Fig. 3b). There was no statistical difference of survival between UAS-Httex1-Q72-eGFP/+;; repo-Gal4/UAS-EDTP-RNAi and UAS-Httex1-Q72-eGFP/+; UAS-EDTP-RNAi/+; repo-Gal4/+ flies. Therefore, downregulation of *EDTP* in glial cells improved the survival with polyQ protein aggregates.

### RNAi knockdown of ***EDTP*** in glial cells or motoneurons extended lifespan

A critical question is whether tissue-specific downregulation of *EDTP* extends lifespan in *Drosophila.*

The lifespan of repo-Gal4/UAS-EDTP-RNAi flies (median 100 days, n = 106) was longer than that of repo-Gal4/+ flies (median 59 days, n = 51) (P < 0.0001, Mantel-Cox test) or that of UAS-EDTP-RNAi/+ flies (median 83 days, n = 78) (Fig. 4a). Therefore, RNAi knockdown of *EDTP* in glial cells extended lifespan in *Drosophila.*

**Figure 4.**
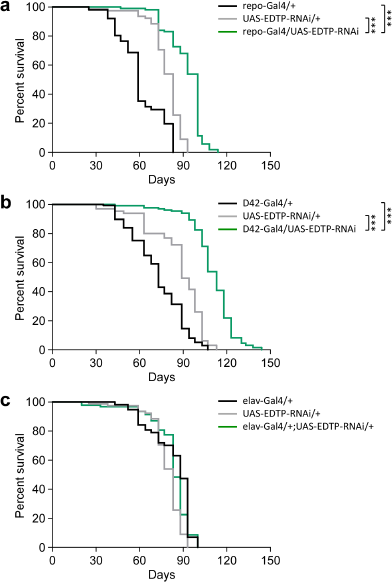
Tissue-specific downregulation of EDTP in glial cells or motoneurons extends lifespan. (a) Lifespan extension by EDTP downregulation in glial cells. Survivorship of repo-Gal4/UAS-EDTP-RNAi (green), repo-Gal4/+ (black) and UAS-EDTP-RNAi/+ (grey) were compared. An EDTP-RNAi line (on the third chromosome) was used. ***, P < 0.0001, Mantel-Cox test. (b) Lifespan extension by EDTP downregulation in motoneurons. Survivorship of D42-Gal4/UAS-EDTP-RNAi (green), D42-Gal4/+ (black) and UAS-EDTP-RNAi/+ (grey) were compared. D42-Gal4 is a motoneuronal driver^22^. ***, P < 0.0001, Mantel-Cox test. (c) Pan-neuronal downregulation of EDTP had no effect on lifespan. elav-Gal4 is a pan-neuronal driver^23^.

Two additional Gal4 drivers, motoneuron-specific D42-Gal4^22^ and pan-neuronal driver elav-Gal4^23^, were used for tissue-specific RNAi knockdown of *EDTP.* The lifespan of D42-Gal4/UAS-EDTP-RNAi flies (median 113 days, n = 132) was extended compared with D42-Gal4/+ flies (median 73 days, n = 137) (P < 0.0001, Mantel-Cox test) or UAS-EDTP-RNAi/+ flies (median 89 days, n = 65) (P < 0.0001, Mantel-Cox test) (Fig. 4b). There was no statistical difference of lifespan between elav-Gal4/+; UAS-EDTP-RNAi/+ flies (median 83 days, n = 93) and elav-Gal4/+ controls (median 88 days, n = 57) or UAS-EDTP-RNAi/+ controls (median 83 days, n = 78) (Fig. 4c). Thus, tissue-specific downregulation of *EDTP* in glial cells or motoneurons extended lifespan in *Drosophila*.

## Discussion

*Drosophila* EDTP is essential in the regulation of oogenesis^5^ as well as muscle performance in adult stage^13^. Downregulation of the mammalian homolog *mJUMPY* promotes the initiation of autophagy^10^. However, loss-of-JUMPY is also associated with muscle dysfunction^14,24^ and a rare genetic disease, centronuclear myopathy^1^. Here we show that tissue-specific downregulation of *EDTP* in non-muscle cells, particularly glial cells, suppresses polyQ protein aggregates expressed in the same cells, improves survival with polyQ protein aggregates, and extends lifespan in *Drosophila*.

It is striking that polyQ protein aggregates can not be detected after around 30 days in flies carrying one EDTP-RNAi construct. Translation of *repo* and expression of repo-Gal4-driven mCD8-GFP in glial cells both persist to at least 56 - 63 days^25^. The RNAi effects must be strong enough to clear accumulated polyQ protein aggregates during young ages (i.e. < 30 days), while at the same time to sufficiently remove newly synthesized aggregates. Several proteins, including histone deacetylase (HDAC)^26^, dHDJ1 (homolog of human HSP40)^27^, dTPR2 (homolog of human tetratricopeptide repeat protein 2)^27^, molecular chaperones HDJ1 and HDJ2^28–30^, human Hsp70^31^, yeast Hsp104^32^, baculoviral antiapoptotic protein P35^33^, and several modifiers for Spinocerebellar ataxia type 1 (SCA1)^34^, are the identified molecular targets for suppressing the expression of polyQ protein aggregates. However, there are no previous reports of complete removal of polyQ protein aggregates by manipulating the expression of these molecules. Therefore, we provide evidence of targeting EDTP for the complete removal of polyQ protein aggregates expressed in glial cells at relatively old ages.

Downregulation of *EDTP* is clearly involved in the suppression of polyQ protein aggregates. By using two different RNAi lines, we observed consistent effects in newly emerged flies and flies around 19 days old, altered expression dynamics of polyQ protein aggregates, and improved survival with polyQ protein aggregates. Additionally, these two RNAi constructs lead to different outcomes. We observed that one RNAi but not another results in disappearance of polyQ protein aggregates, and that two RNAi lines have different modifications on the expression dynamics of polyQ protein aggregates. Each RNAi line carries a short hairpin RNA (shRNA) targeting a unique exon of *EDTP.* A specific *EDTP* exon could have specific splicing and abundance characteristics. This could explain the different suppression of aggregates between two independent RNAi lines. Furthermore, the shRNAs in two lines each have potential off-target effects. The shRNA in the RNAi line which results in complete disappearance of aggregates has a potential target gene *Traf-like* (CG4394), encoding TNF-receptor-associated factor-like. Traf-like is inferred to have ubiquitin-protein transferase activity (http://flybase.org/reports/FBgn0030748.html), which might be harnessed by downregulation to assist in the suppression of polyQ protein aggregates. The shRNA in the second

RNAi line has two potential off-targets, *CG5142* and *ORCT.* The product of *CG5142* has a tetratricopeptide repeat domain (http://flybase.org/reports/FBgn0032470.html). A *Drosophila* tetratricopeptide repeat protein dTPR2 has been shown to suppress polyQ protein aggregates in the compound eyes^27^. *ORCT* encodes an organic cation transporter (http://flybase.org/reports/FBgn0019952.html). How ORCT affects the removal of polyQ protein aggregates is currently unclear. Downregulation of potential off-targets might modify the effects of RNAi knockdown of *EDTP,* resulting in different suppression of polyQ protein aggregates.

Both RNAi lines improves the maximal survival with polyQ protein aggregates to a similar level (around 2 fold). Thus, differential suppression of polyQ protein aggregates has little effect on the improvement of survival. The expression of polyQ protein aggregates greatly restrict the maximal lifespan to around 20 days. A doubling of survival is remarkable, but might have reached the maximal limit of the effect of EDTP downregulation. Perhaps, molecular candidates that lead to further suppression of polyQ protein aggregates could have a greater role than EDTP in the improvement of survival. Comparison between the improved survival (with the expression of polyQ protein aggregates) to the lifespan that does not involve polyQ protein aggregates is less meaningful, unless one has identified a molecular target that is responsible for complete removal of polyQ protein aggregates throughout the lifespan.

In addition to the suppression of polyQ protein aggregates, lifespan extension through *EDTP* downregulation in glial cells firmly supports the hypothesis that tissue-specific manipulation of EDTP expression circumvents the mutation-related muscle phenotypes, shifting the overall effect to lifespan extension. A *Drosophila* heterozygous *EDTP* mutant appears to have unaffected muscle function and lifespan^13^. It suggests that one genomic copy of *EDTP* is sufficient to produce normal level of EDTP in muscles. However, heterozygous *EDTP* mutation might cause reduced EDTP in non-muscle tissues, which would affect the balance of PtdIns3P and promote autophagy. This is supported by the findings that heterozygous *EDTP* flies have improved survival after prolonged anoxia, a condition inducing autophagy in mammalian and human cells^17^. Through tissue-specific downregulation of *EDTP,* we also find that motoneurons are another target tissue favored for lifespan extension. Motoneuronal overexpression of a human CuZn superoxide dismutase (SOD1) extends *Drosophila* lifespan^22^. By targeting pan-neurons in the central nervous system for *EDTP* downregulation, we observe no beneficial effect in lifespan extension. This would be because of possibly low expression of EDTP in targeted neurons, or these cells have developed processes independent of EDTP for self clearance/disposal of protein aggregates.

EDTP/JUMPY represents a novel phosphoinositide phosphatase that belongs to myotubularian (MTM) family^1^. EDTP/JUMPY hydrolyzes 3-position phosphate from PtdIns3P and PtdIns(3,5)P_2_. Mouse JUMPY negatively controls PtdIns3P levels^10^. PtdInd3P regulates many aspects of autophagy. In the early events of autophagy, PtdInd3P recruits effectors and initiates the formation of autophagosome^35^. Later, PtdInd3P attracts MTM3 for its turnover and promotes autophagosome maturation into autolysosome^36^. Thus, JUMPY is actively involved in the regulation of autophagy. In addition, *JUMPY* mutation causes the accumulation of PtdIns(3,5)P_2_ in skeletal muscles. Accumulated PtdIns(3,5)P_2_ binds directly to ryanodine receptors, RyR1 and RyR2, in sarcoplasmic reticulum and increases intracellular Ca^2+^ in both skeletal and cardiac muscle^14,24^, resulting in muscle dysfunction^14^ and altered cardiac contractility^24^.

Abundant expression of EDTP/JUMPY in muscles could be a barrier for utilizing the effect of negative regulation of autophagy through ubiquitous downregulation. Tissue-specific manipulation of EDTP expression by targeting non-muscle tissues circumvents this problem. The advantage of tissue-specific downregulation of EDTP is to avoid or minimize the *EDTP*/*JUMPY*-mutation-associated muscle phenotypes. Targeting glial cells provides an additional advantage. Restricted downregulation of EDTP in glial cells causes no or minor disturbance to maternally derived EDTP expression in the development of oocytes. Transcription of *repo,* which is trapped for Gal4 expression in repo-Gal4 fly, first appears to be highly restricted in glioblasts in stage 9 embryos^19^. *EDTP* mRNA, however, is detected uniformly in the cytoplasm of eggs at stage 1, 5 and 11 but mostly disappears at stage 15 and after^5^. There is little spatial and temporal overlap of transcription between *repo* and *EDTP* during oogenesis. Therefore, glial cell-specific downregulation of EDTP occurs mostly at adult stage, during which EDTP reappears around day 7, peaks at day 20-30 and subsequently decreases in whole-fly preparations^6^.

In conclusion, we demonstrate an approach to fine-tune the expression of a disease-associate gene *EDTP/JUMPY* for the suppression of polyQ protein aggregates and lifespan extension in *Drosophila*.

## Methods

### Flies

Fly strains and their sources are: DJ694^6^, repo-Gal4 (Bloomington *Drosophila* Stock Center (BDSC) #7415); D42-Gal4 (BDSC #8816); elav-Gal4 (BDSC #8765); UAS-EDTP-RNAi (with an RNAi construct on the second chromosome, BDSC #41633); UAS-EDTP-RNAi (with an RNAi construct on the third chromosome, BDSC #36917); UAS-Httex1-Q72-eGFP^18^; UAS-hsp70-myc^21^; 20×UAS-IVS-mCD8::GFP (BDSC #32914) and w^1118^. We recombined several transgenes and generated these flies: UAS-EDTP-RNAi (on II); UAS-EDTP-RNAi (on III) and UAS-Httex1-Q72-eGFP;;repo-Gal4/TM3. Flies were maintained with standard medium (cornmeal, agar, molasses and yeast) at 21-23 °C in a 12/12 h light/dark condition. Male flies were used for the experiments, unless otherwise stated.

Heterozygous DJ694 flies and their sibling controls were prepared by two consecutive crosses between homozygous DJ694 and w^1118^ flies. We chose virgin female progenies from first mating between male DJ694 and female w^1118^, and crossed them to w^1118^ males. Flies carrying EDTP mutation (red eyed) and their siblings (white eyed) were collected for the tests of survival after prolonged anoxia.

### Survival after prolonged anoxia

Flies were exposed to a 6-h anoxia (generated by pure nitrogen gas) at the ages of 7-8 days. Dead flies were scored daily during the first week of recovery. Thereafter, dead flies were scored twice a week until all the flies were counted. Throughout the experiments alive flies were transferred to fresh food vials twice a week.

### Immunohistochemistry

Immunohistochemistry was performed by following two similar protocols^37,38^. Briefly, dissected brains were fixed in freshly prepared 4*%* paraformaldehyde for 1 h. After three washes with PAT (PBS with 0.5 *%* BSA and 0.5 *%* Triton X-100), tissues were incubated with saturation buffer (10 *%* goat serum in PAT) for 1 h at room temperature. Brain tissues were then incubated with primary antibody at 4 °C for 1-2 days with gentle rotation. Following three washes, tissues were incubated with appropriate secondary antibody. Incubation with secondary antibody was performed at 4 °C in a dark room for 1-2 days with gentle rotation. Flies and their specific antibodies were: (1) repo-Gal4/UAS-hsp70-myc flies, primary antibody: rabbit anti-cmyc (A00173, GenScript) at 1:50, secondary antibody: DyLight goat anti-rabbit IgG (111-485-144, Jackson ImmunoResearch) at 1:500. (2) repo-Gal4/20xUAS-IVS-mCD8::GFP flies, primary antibody: mouse anti-GFP supernatant (12A6, DSHB) at 1:20, secondary antibody: Alexa Fluor 488 conjugated goat anti-mouse IgG (115-545-003, Jackson ImmunoResearch) at 1:500. After three washes tissues were suspended in 200 μl SlowFade Gold antifade reagent (S36938, Life Technologies) and mounted on slides for microscopy. Image stacks were taken using a Carl Zeiss LSM 710NLO laser scanning confocal/multiphoton microscope (Carl Zeiss), and processed with ImageJ (NIH).

PolyQ-expressing flies were imaged without immunohistochemistry. Briefly, fly brains with polyQ expression were dissected in Schneider’s insect medium (S0146, Sigma-Aldrich) containing 0.5 *%* Triton-X-100, mounted on a slide immediately with a small drop of SlowFade Gold antifade reagent (S36938, Life Technologies), and imaged within an hour. The images of Z stacks spaced by 5-7*μ*m throughout the thickness of tissue were taken.

### Quantification of polyQ protein aggregates

Relative expression of polyQ protein aggregates in the brain was evaluated by calculating the percent area for polyQ protein aggregates in the brain (*%* PolyQ area). Image stacks were combined to a single plane of image file. The total area of aggregate particles was measured using ImageJ software. *%* PolyQ area was calculated by this formula: *%* PolyQ area = Total area of aggregate particles/The area of brain × 100 *%*. For each genotype of flies, at every time point two samples were used for evaluating the expression of polyQ protein aggregates.

### Lifespan experiments

Flies were collected at 0-2 days after emergence at a density of 20-25 flies per vial. They were transferred into fresh food vials twice a week. Dead flies were scored during transfer until all the flies were counted. This procedure was also used for the examination of the lifespan of flies expressing ployQ protein aggregates in glial cells, and the lifespan of flies having simultaneous expression of ployQ protein aggregates and RNAi knockdown of EDTP in glial cells. The lifespan experiments were repeated once. Sample sizes are indicated in the text.

### Statistics

Mantel-Cox tests were performed for survival analysis. Two-way ANOVA with Bonferroni post-tests were used for comparing relative expression of polyQ protein aggregates between different flies. Statistics were performed using GraphPad Prism5 software. A *P* < 0.05 was considered a significant difference.

## Acknowledgements

This work was funded by Natural Sciences and Engineering Research Council of Canada (NSERC) grant (RGPIN 40930-09) to R.M.R.

## Author contributions statement

C.X. contributed to experimental design; C.X. and S.Q. contributed to data collection, analysis and manuscript preparation; R.M.R. contributed to funding support and manuscript editing; L.S contributed to research materials, comments and discussion. All authors reviewed the manuscript.

## Additional information

The authors declare no conflict of interest.

## Supplementary Information

**Figure S1.**
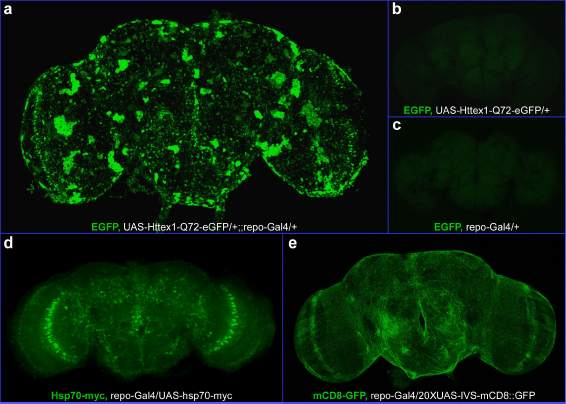
PolyQ protein aggregates expressed in glial cells. (a) PolyQ protein aggregates expressed in glial cells in UAS-Httex1-Q72-eGFP/+;;repo-Gal4/+ fly. Immunostaining of the target proteins (in green) and fly genotype (in white) are indicated. Large protein aggregates are seen located around the surface of the brain. (b) No detectable protein aggregates in UAS-Httex1-Q72-eGFP/+ fly. (c) No detectable protein aggregates in repo-Gal4/+ fly. (d) Expression of a soluble molecular chaperone (Hsp70-myc) in glial cells in repo-Gal4/UAS-hsp70-myc fly. Experiment was repeated here from a previous report^39^ as a comparison for the expression of polyQ protein aggregates. (e) Expression of a membrane protein (mCD8-GFP) in glial cells in repo-Gal4/20×UAS-IVS-mCD8::GFP fly. Male flies at 1-7 days were used for imaging.

**Figure S2.**
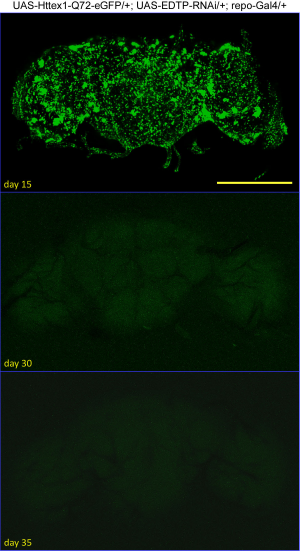
Suppression of polyQ protein aggregates in glial cells by ***EDTP*** downregulation. Expression of polyQ protein aggregates were invisible at day 30 and 35 in UAS-Httex1-Q72-eGFP/+; UAS-EDTP-RNAi/+; repo-Gal4/+ flies. Shown are the brains of males at 15, 30 and 35 days. Male flies are used for imaging. Bar 200 *μ*m.

**Figure S3.**
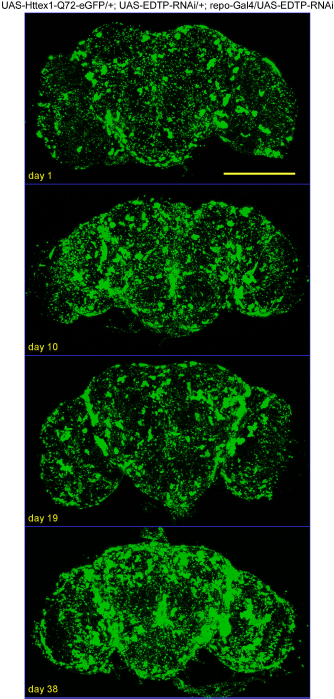
Expression of polyQ protein aggregates in glial cells with double *EDTP*-RNAi. Expression of polyQ protein aggregates and simultaneous knockdown of ***EDTP*** by double RNAi in glial cells. Fly genotype: UAS-Httex1-Q72-eGFP/+; UAS-EDTP–RNAi/+; repo-Gal4/UAS-EDTP-RNAi. Shown are the brains of males at 1,10,19 and 38 days. Male flies are used for imaging. Bar 200 *μ*m.

